# A Novel Mice Model of Catecholaminergic Polymorphic Ventricular Tachycardia Generated by CRISPR/Cas9

**DOI:** 10.1101/2021.10.14.464343

**Authors:** Cuilan Hou, Xunwei Jiang, Qingzhu Qiu, Junmin Zheng, Shujia Lin, Shun Chen, Meng Xu, Yongwei Zhang, Lijian Xie, Tingting Xiao

## Abstract

Catecholaminergic polymorphic ventricular tachycardia (CPVT) has been considered as one of the most important causes of children’s sudden cardiac death. Mutations in the genes for RyR2 and CASQ2, two mainly subtypes of CPVT, have been identified. However, the mutation in the gene of TECRL was rarely reported, which could be another genetic cause of CPVT. We evaluated myocardial contractility, electrophysiology, calcium handling in Tecrl knockout (Tecrl KO) mice and human induced pluripotent stem cell-derived cardiomyocytes. Immediately after epinephrine plus caffeine injection, Tecrl KO mice showed much more multiple premature ventricular beats and ventricular tachycardia. The Tecrl KO mice demonstrate CPVT phenotypes. Mechanistically, intracellular calcium amplitude was reduced, while time to baseline of 50 was increased in acute isolated cardiomyocytes. RyR2 protein levels decreased significantly upon cycloheximide treatment in TECRL deficiency cardiomyocytes. Overexpression of TECRL and KN93 can partially reverse cardiomyocytes calcium dysfunction, and this is p-CaMKII/CaMKII dependent. Therefore, a new CPVT mouse model was constructed. We propose a previously unrecognized mechanism of TECRL and provide support for the therapeutic targeting of TECRL in treating CPVT.

## 1. Introduction

CPVT is a life-threatening inherited arrhythmia characterized by stress- or emotion-induced bipolar ventricular preexcitations and polymorphic ventricular tachycardias (VT)(Hou et al, 2021). Since 1970, several CPVT related gene mutations have been found(van der Werf & Wilde, 2013). Priori’s group reported that cardiac ryanodine receptor mutations caused CPVT and approximately 50% of CPVT patients carried mutations in the ryanodine receptor 2 (RyR2)(Hayashi et al, 2009; Priori et al, 2001). Lahat *et al* showed that cardiac calsequestrin gene (CASQ2) mutations are responsible for the rare autosomal recessive form, which accounts for approximately 3%-5% of CPVT(Lahat et al, 2001). In addition, there are other autosomal recessive CPVT subtypes, including the integral membrane protein encoding gene triadin (TRDN)(van der Werf & Wilde, 2013) or trans-2, 3-enoyl-CoA reductase-like (TECRL), which was recently suggested to be a novel candidate gene for life-threatening inherited arrhythmias (Moscu-Gregor et al, 2020).

Despite the facts that the profound research of genetic loci of CPVT have been approached over the last decade, the understanding of other phenotypes of CPVT remains challenging (Schwartz et al, 2013). In several CPVT probands, TECRL mutations have been identified to be pathogenic, which dawned on us that this may represent a novel clinical phenotype of CPVT. *Via* whole-exon sequencing (WES), an Arg196Gln was detected in 2 French Canadian patients and c.331+1G>A of TECRL was revealed in affected members of a Sudanese family(Devalla et al, 2016). An Arg196Gln and c.918+3T > G splice site mutation in *TECRL* gene was firstly reported in a Chinese family (Xie et al, 2019). Alexander Moscu-Gregor’s group recently summarized three TECRL mutations among 631 index patients with suspected CPVT or LQTS, which are homozygous for Gln139*, homozygous for Pro290His, compound heterozygous Ser309*, and Val298Ala (Moscu-Gregor et al, 2020). The above findings suggested that several *TECRL* mutations were identified with CPVT (Devalla et al, 2016; Moscu-Gregor et al, 2020; Xie et al, 2019), but the protein function and pathogenic mechanism are still needed to further be elucidated.

It was well known that RyR2 and CASQ2 gene mutations induce the release of Ca^2+^ from the sarcoplasmic reticulum (SR) during diastolic period, leading to intracellular calcium overload (Knollmann et al, 2006). The increase of Ca^2 +^ content in the cell activates the activity of Na^+^-Ca^2+^-exchanger (NCX), which produces transient inward current. The delayed depolarization of calcium transient inward current lead to delayed after depolarizations (DADs), further inducing VT and fibrillation (Bezzerides et al, 2019). RyR2 is the release channel of SR calcium, which can increase the release of calcium in SR during diastolic period (Kurtzwald-Josefson et al, 2012; Manotheepan et al, 2016). CASQ2 gene mutation causes the decrease of SR calcium buffering capacity and increase of intracellular calcium concentration (Kurtzwald-Josefson et al, 2012). The imbalance of calcium homeostasis in cardiomyocytes is considered as an important mechanism of ventricular arrhythmias (Park et al, 2019). In 2016, Devalla’s group first reported that lower amplitude of diastolic Ca^2+^ transient and slower decay of cytosolic Ca^2+^ in the homozygous cardiomyocytes derived from the CPVT patient hiPSCs (Devalla et al, 2016). However, the *TECRL* protein functions in CPVT or other cardiovascular disease are still not well understood. Neither does the mechanisms linking *TECRL* deficiency with CPVT. *TECRL* may be a novel CPVT disease-causing gene. To clarify the molecular mechanism of *TECRL* induced CPVT is important to provide crucial clues for the diagnosis and treatment of CPVT.

In this study, we developed a Tecrl knockout (Tecrl KO) mice model via CRISPR/Cas9 technique to decipher the role of *TECRL* dysfunction in the context of CPVT pathobiology, which represents an interesting model to study its related cellular and molecular mechanisms. Using a combination of electrophysiological approaches, histological analyses, and molecular biology, we characterized the first Tecrl KO mice model and provided tools for developing specific therapeutic targets for CPVT. Our findings shed light on the complex pathogenesis of recessive CPVT.

## 2. Results

### 2.1 Generation of the Tecrl KO mice using CRISPR/Cas9 and its effects on cardiac pathology

In order to disrupt the open reading frame of Tecrl in mice, we designed a pair of sgRNAs targeting mice Tecrl (Fig. 1A). The sequences of sgRNAs were shown in Table s1. The genotypic of Tecrl KO mice were confirmed by PCR amplification and DNA sequencing. Primers for genotype identification and RT-PCR were shown in Table 1. RT-PCR confirmed the loss of Tecrl, and we found Ryr2 was also decreased significantly in the heart of Tecrl KO mice compared with that of the wild type (WT) (Fig. s1 A-B). We tested the blood samples from a CPVT patient who has a TECRL compound heterozygous mutation (Arg196Gln and c.918+3T > G) and his parents (Xie et al, 2019), and found that TECRL protein level in the patient was much lower than his parents’ (Fig. s1 C). No differences were observed in the duration of the pregnancy, delivery, size, and survival of litters between the WT and Tecrl KO mice.

**Table 1.**
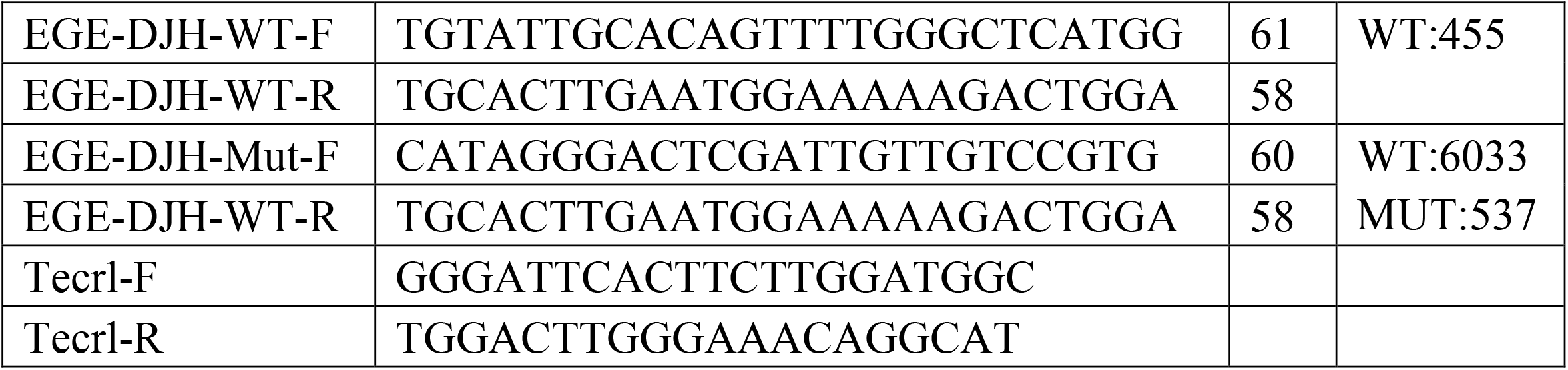
Primers sequence of genotype identification and RT-PCR.

**Fig 1.**
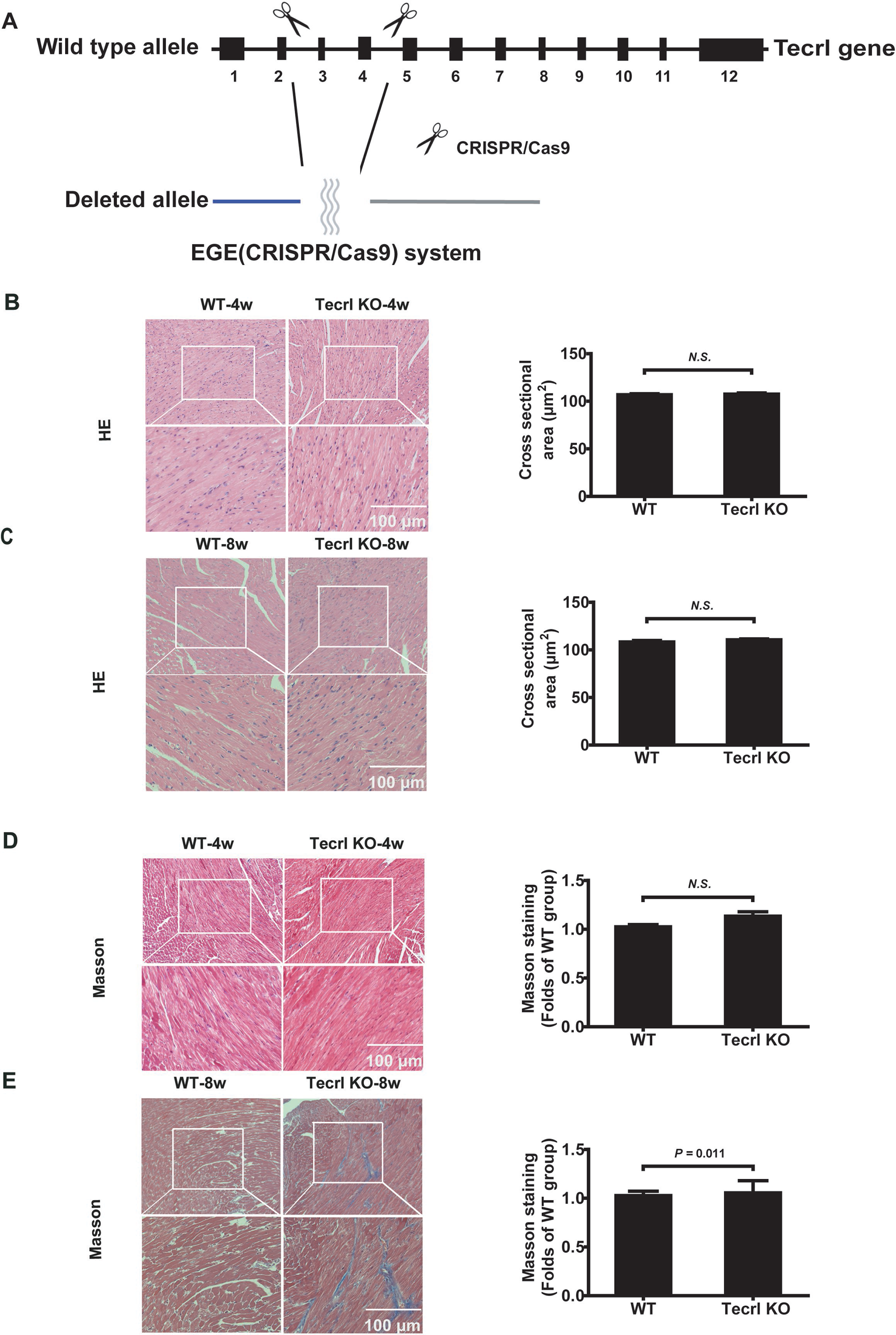
Generation of Tecrl KO mice using CRISPR/Cas9 and its effects on cardiac pathological state. A. The general structure of Tecrl gene and the schematic diagram of two sgRNA target sites located in exon 3 and exon 4 of the mice Tecrl locus. B-C. Representative images of HE and quantification of cross sectional area in four- and eight-week mice (WT group, n=4; Tecrl KO group, n=6). D-E. Representative images of Masson staining and quantification of fibrosis percentages in four- and eight-week mice (WT group, n=4; Tecrl KO group, n=7). Values are means ± SE. *P* < 0.05 was considered significant.

To examine whether Tecrl deficiency in mice causes any pathology in the heart, we performed histological assessments of these mice. Compared with the WT mice, the Tecrl KO mice cardiomyocytes exhibited similar in size both at the age of four and eight weeks (Fig. 1B-C). There is no significance of interstitial collagen volume between the WT and Tecrl KO mice at the age of four weeks (Fig. 1D). However, the Tecrl KO mice exhibited some pathological changes, such as increased interstitial collagen volumes, compared with the WT mice at the age of eight weeks (Fig. 1E).

We observed no significant difference in cell death (TUNEL-positive myocytes) between the WT and Tecrl KO at the age of four weeks (Fig. 2A-B). There were much more TUNEL-positive myocytes in the Tecrl KO mice than the WT at the age of eight weeks (Fig. 2C-D). And this trend of increased apoptosis was also confirmed by increased Bax protein levels and decreased Bcl-2 levels (Fig. 2E-F).

**Fig 2.**
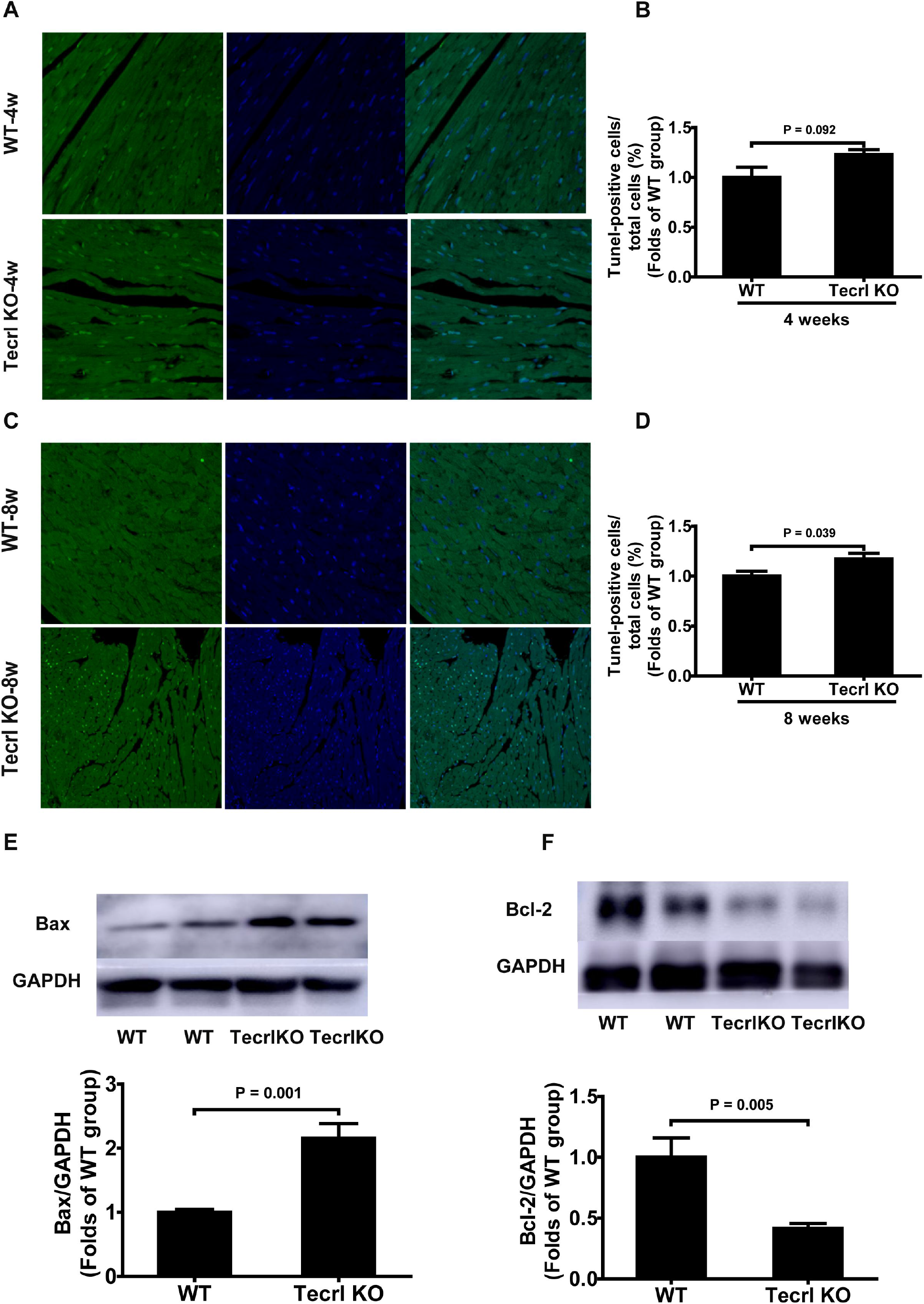
Tecrl deficiency effects on cardiac apoptosis. A-B. Myocardial apoptosis was determined and quantified by TUNEL assay in four weeks mice. C-D. Myocardial apoptosis was determined and quantified by TUNEL assay in eight weeks mice. The arrow showed TUNEL-positive cells; nuclei: the blue; scale bar = 20 μm. E-F. Representative of western blotting images and quantification of the expression Bax and Bcl-2 in eight weeks mice (n=6). Values are means ± SE. *P* < 0.05 was considered significant.

All of the above data indicate that long time Tecrl deficiency results in myocardial pathology. Therefore, subsequent experiments were conducted in eight weeks animals.

### 2.2 Evaluation the incidence of ventricular arrhythmias in adult Tecrl KO mice

None of the mice demonstrated ventricular events at resting condition prior to drug administration. In the Tecrl KO mice, multiple premature ventricular beats and ventricular tachycardia (VT) were observed immediately after epinephrine plus caffeine (epi/caffeine) injection; however, few ventricular events were observed in the WT mice (Fig. 3A-B). Both bigeminy and sustained VT were observed in the Tecrl KO mice rather than the WT. And the Tecrl KO mice VT episode (e.g. one to five seconds, five to fifteen seconds, or above fifteen seconds)sustained much longer than the WT (Fig. 3C-D). Eight-week-old Tecrl KO mice PR interphase was considerably shorter than the WT mice after epi/caffeine stimulation. Both QRS and QT interphases were noticeably longer in the 8-week-old Tecrl KO mice than the WT (Fig. 3E).

**Fig 3.**
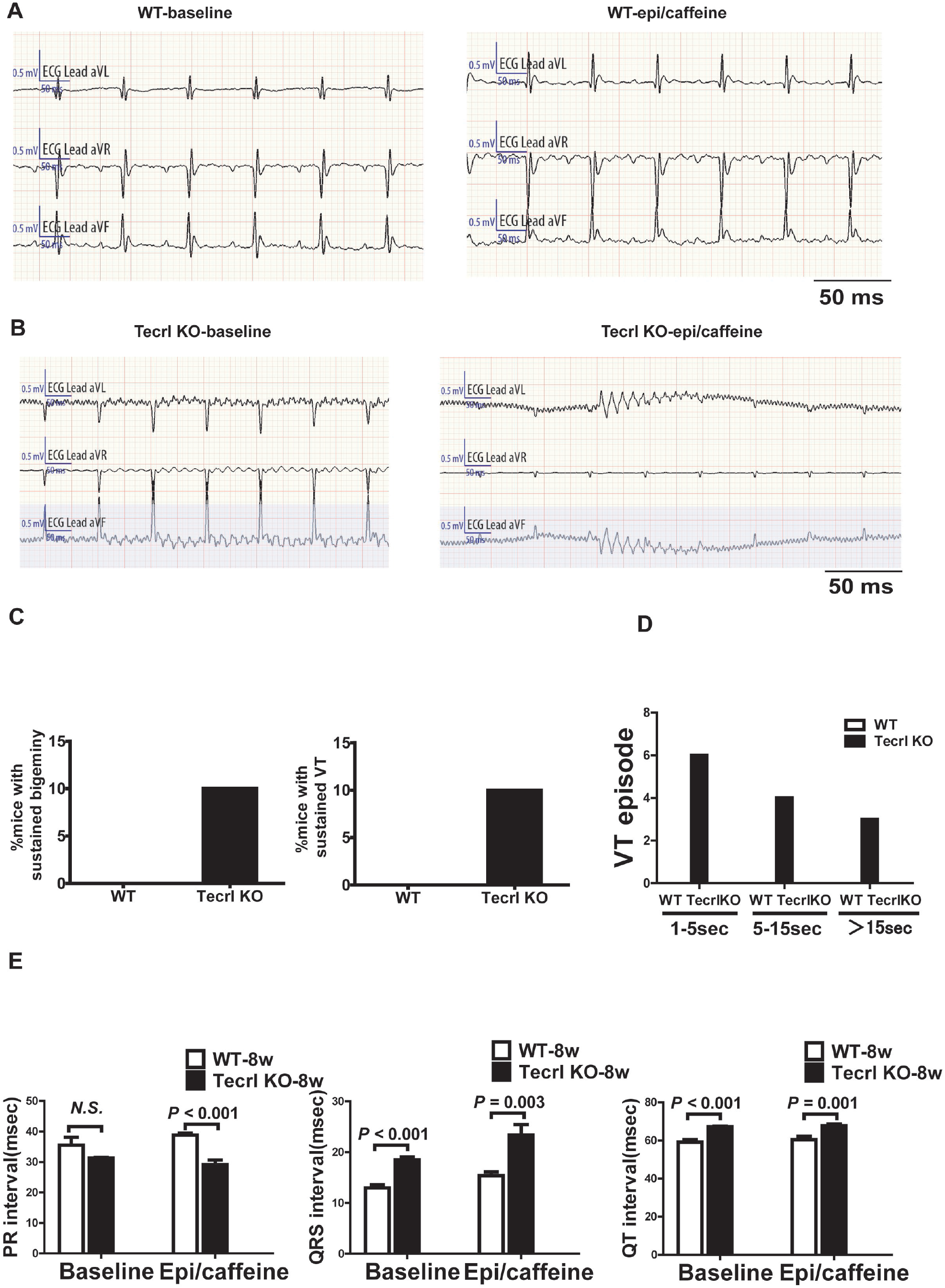
Arrhythmia inductions in the Tecrl KO mice of eight weeks. A-B. Representative ECG recordings in the WT and Tecrl KO mice before and after epi/caffeine stimulation. C-D. Quantification of the sustained bigeminy, sustained VT, and VT episode in the WT and Tecrl KO mice. E. Quantification of the PR, QRS, and QT in the WT and Tecrl KO mice (n=10). Values are means ± SE. *P* < 0.05 was considered significant.

CPVT is characterized by ventricular tachycardia, or syncope due to emotions or physical exercise and without structural heart defects(Hayashi et al, 2009). And CPVT is part of the prevalent causes of sudden cardiac death during childhood and in adolescence(Hayashi et al, 2009). We tested 4/5 weeks heart histological and functional assessments. Similarly, the 4/5 weeks Tecrl KO mice were more prone to CPVT phenotypes induced by epi/caffeine. Briefly, multiple premature ventricular beats and VT were observed immediately after epi/caffeine injection, whereas few ventricular events were observed in the WT mice (Fig. s2 A-B). There was not any significance of PR interphase between the WT and Tecrl KO mice. Both QRS and QT interphases were significantly shorter in the 4/5-week-old Tecrl KO mice than the WT (Fig. s2 C). Thus, the Tecrl KO mice demonstrate CPVT phenotypes.

### 2.3 Cell shorting

Next, we studied intracellular calcium transients of four to six independent mice cardiomyocytes isolation in the WT and Tecrl KO mice. The measures of sarcomere shortening were recorded in the presence of 5 μmol/L Fura-2 AM and stimulated at 1 Hz. A summary of the canine cardiomyocytes in the WT and Tecrl KO mice characteristics is presented in Fig. 4A-B. The Tecrl KO mice showed some abnormities in the heart mitochondria, such as irregular arrangement, swelling, vacuolated and disrupted cristae (Fig. 4A). The cTnT staining, one of the cardiomyocyte’s marker, demonstrates that the high quality of the acute isolated cardiomyocytes (Fig. 4B). The calcium transient increase is markedly slower in the Tecrl KO mice than the WT (Fig. 4C), resulting in a significant difference in the time required to reach 50% of the calcium transient amplitude (peak and baseline) (Fig. 4D, 4H). The decay rate of the caffeine-induced Ca^2+^ transient showed no difference between the WT and Tecrl KO mice cardiomyocytes (Fig. 4E-F). The time required to reach peak 50% was decreased in the Tecrl KO mice compared with the WT, while there were no significant differences in these two groups (Fig. 4G).

**Fig 4.**
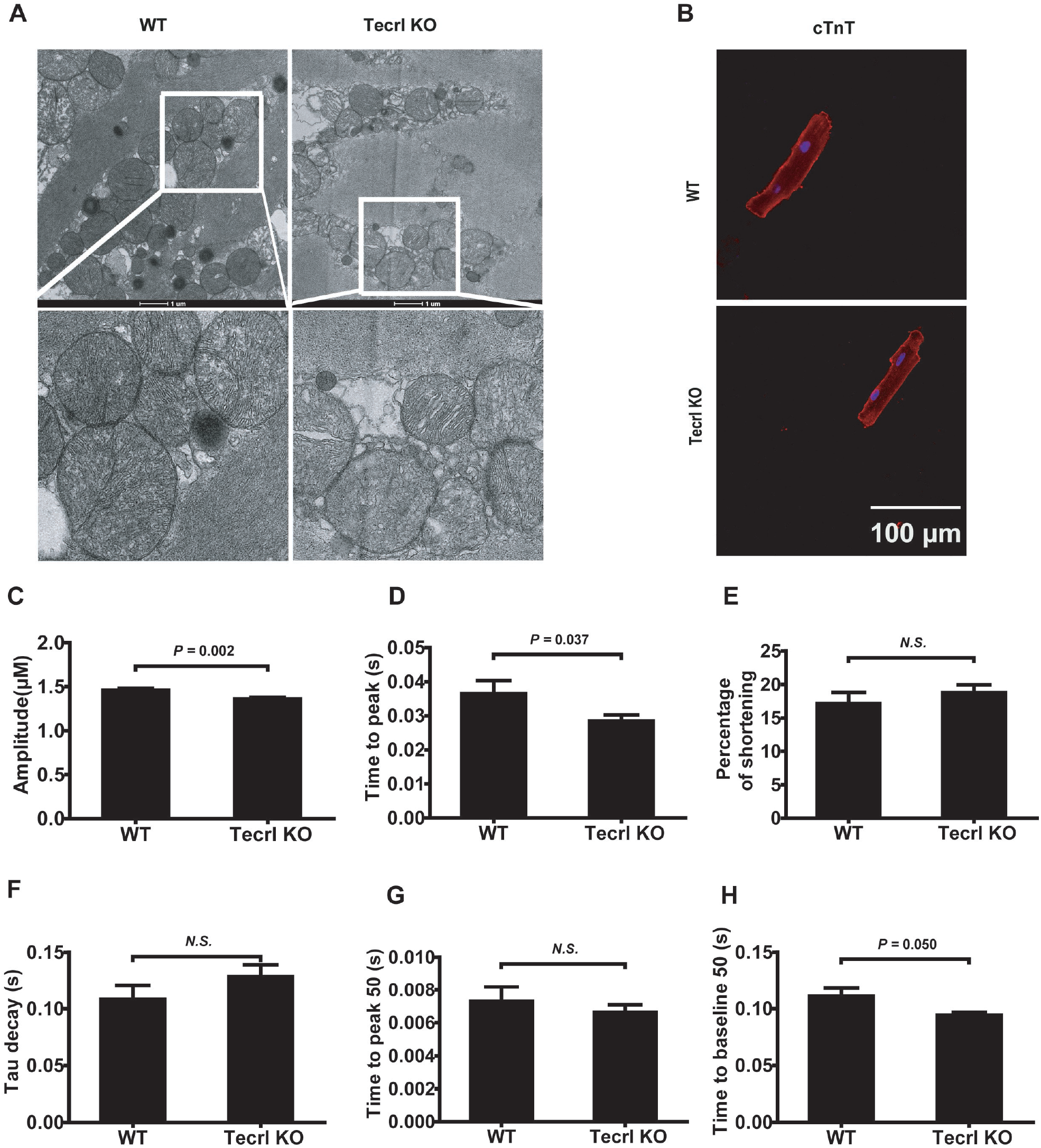
Tecrl deficiency effects on cardiomyocytes calcium handling. A. Cardiac ultrastructure of the cardiomyocytes was examined by TEM. Representative images of TEM in acute isolated mice cardiomyocytes (n=3). Magnification is 6700, scale bar=1 μm. B. Representative immunofluorescence images of cTnT in the acute isolated mice cardiomyocytes (n=6). C-H. Representative calcium handling images and quantification of the amplitude (C), time to peak (D), percentage of shorting (E), Tau decay (F), time to peak 50 (G),time to baseline 50 (H) (11 cells from four WT mice, 28 cells from five Tecrl KO mice). Values are means ± SE. *P* < 0.05 was considered significant.

### 2.4 Global impacts of Tecrl KO on cardiac gene expression

To determine the impact of disrupted Tecrl signaling on cardiac gene expression, transcription analysis was performed in the Tecrl KO mice ventricular muscle in comparison with age-matched litter mate controls. We identified a robust signature of mRNA within cardiac tissues (181) isolated from the WT and Tecrl KO mice. To understand the signaling pathways most affected by deficiency of Tecrl, GO enrichment analysis of differentially-expressed mRNAs was performed to reveal disproportionate representation of mRNA involved in heart contraction, cytosolic calcium ion concentration, glucose metabolism, and muscle contraction (Fig. 5A). Then we measured and confirmed the related calcium signaling pathways using western blotting assay (Fig. 5B-G). We examined the expression levels of CaMKII in the mouse cardiomyocytes. Tecrl knockout significantly activated CaMKII phosphorylation in the mice hearts (Fig. 5B-C). Previous study showed that CaMKII modulates glucose metabolism and calcium signaling through its downstream mediator P38 mitogen activated protein kinase, and P38 is also involved in calcium regulation(Ozcan et al, 2012). We also found that Tecrl knockout also activated p38 phosphorylation in mice heart (Fig. 5B, 5D). A series of calcium regulating signaling, such as Casq2 or Ryr2, were decreased in the Tecrl KO mice heart than that of the WT (Fig. 5B, 5E-F). Expression levels of Ncx still remained the same between the WT and Tecrl KO mice (Fig. 5B, 5G). Then we used an *in vitro* translation inhibition study to test the relationship between TECRL and RyR2, and we found that after 4 h of treatment with CHX, the RyR2 expression level of the shTECRL group was significantly reduced. However, the degradation of the Cth group was considerably slower, and the protein bands were still visible even after 24 h, indicating that TECRL deficiency could accelerate RyR2 protein degradation *in vitro* (Fig. s3A-B).

**Fig 5.**
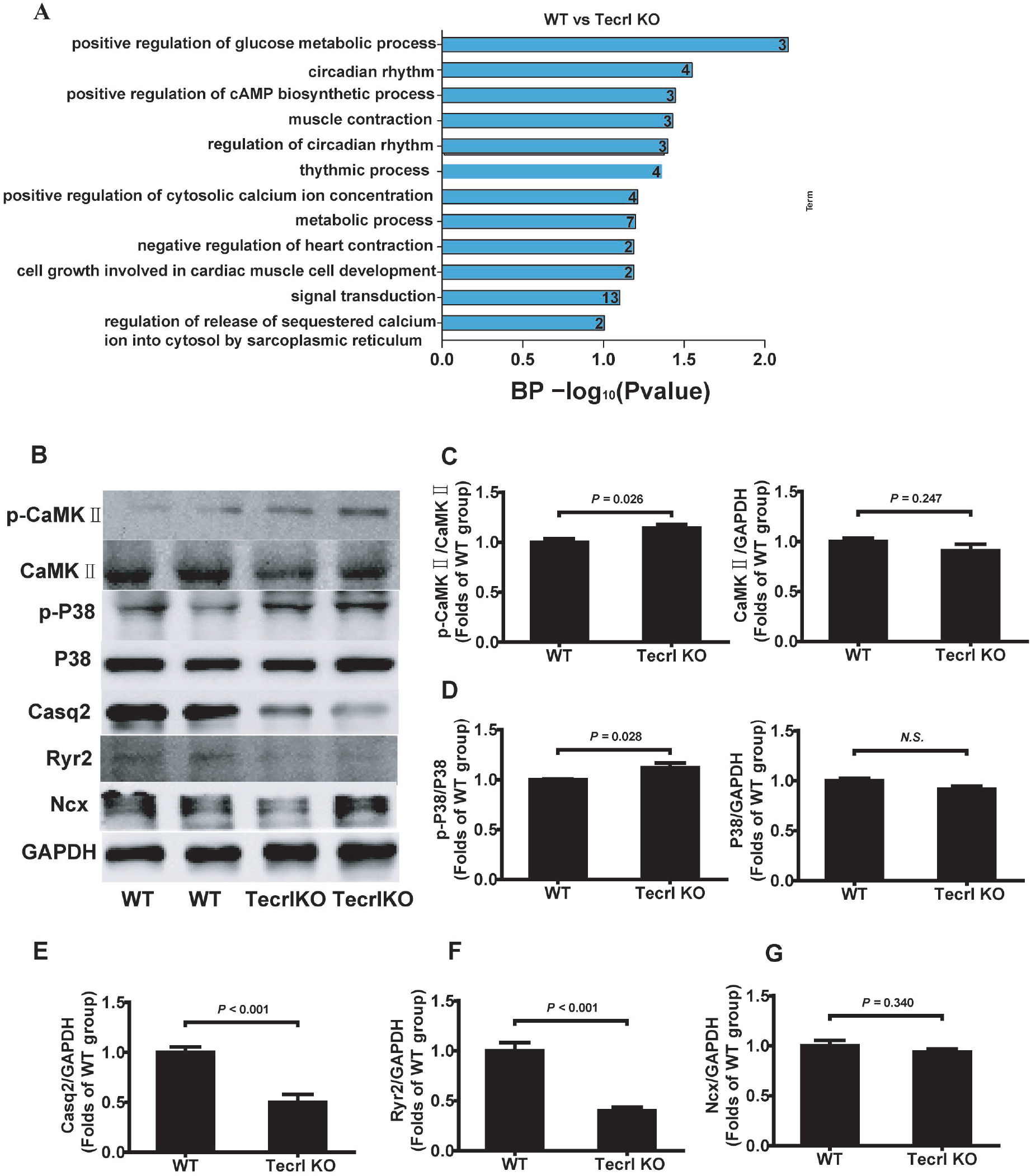
Visualization of differentially regulated mRNAs in cardiac tissues of the WT and Tecrl KO mice. A. GO analyses of differentially-expressed mRNAs. B-G. Representative of western blotting images and quantification of p-P38, P38, Casq2, Ncx, Ryr2, p-CaMKII, and CaMKII expression in the WT and Tecrl KO mice (n=6). Values are means ± SE. *P* < 0.05 was considered significant.

### 2.5 Contraction of hiPS-CMs

To demonstrate the stability of our cell model based on our previously established protocol(Hou et al, 2021), we first stained SOX2 and OCT4. HiPS-CMs are highly expressed in SOX2 and OCT4, showed that these cells maintained stem cells characteristics of undifferentiated stem cells (Fig. 6A). Then these stem cells were directional differentiation into cardiomyocytes using established protocol(Chang et al, 2016). Fig. 6B showed that these cells highly expressed in cTnT, indicating that these cells were hiPSC-CMs. At the same time, we found that TECRL was colocalized with CALNEXIN, an endoplasmic reticulum chaperone. Furthermore, we measured the baseline contraction of hiPSC-CMs. Statistical analysis showed no significant difference between the Cth and shTECRL groups (Table 2). Pacing at 1 Hz with 5V for 300 ms, cardiomyocytes were able to be syncronized at the preliminary rate. Velocity/pixel/sec of the above two groups showed no significant difference (Table 3), confirming the possibility to measure the contraction of hiPSC-CMs. Positive intrope pharmacological effect of hiPSC-CMs was tested. Isoprenaline (ISO, 1µM) induced faster beating rate of TECRL knockdown hiPSC-CMs versus baseline (Table 4). Representative absolute amplitude and rhythm of hiPSC-CMs are shown in (Fig. s3). ISO evoked significantly higher contraction peak value of TECRL knockdown hiPSC-CMs compared to baseline (Fig. s4B, 4D), while ISO had no statistically significant effect in control hiPSC-CMs (Fig. s4A, 4C). At the same time, we overexpressed TECRL, and found that it did not alter the cardiomyocytes contraction (Table 5) and velocity (Table 6). ISO evoked significantly higher contraction peak value of TECRL overexpressed hiPSC-CMs compared to baseline (Table 7). Next, we used hiPSC-CMs with a commercially available 96-well screening platform (CardioExcyte 96, Nanion Technologies GmbH) for impedance measurements. Under controlled conditions, cardiomyocytes displayed spontaneously beating and impedance signals with a beat rate of 55.95 beats per minute. Under ISO stimulation, cardiomyocytes isolated from Tecrl KO mice were beating faster, their base impedance was increased, while amplitude kept stable compared with the WT cardiomyocytes (Fig. 6C-F).

**Table 2.** Baseline of contraction of TECRL knockdown hiPSC-CMs versus baseline.

|  | n | bpm (mean±SD) | Rate (Hz) | *p* | *t* |
| --- | --- | --- | --- | --- | --- |
| Cth | 11 | 37.6 ± 19.85 | 0.63 | 0.4639 | 0.7419 |
| shTECRL | 17 | 33.2 ± 12.33 | 0.55 |  |  |

**Table 3.** Velocity/pixel/sec of TECRL knockdown hiPSC-CMs versus baseline.

|  | n | Velocity/pixel/sec (mean±SD) | *p* | *t* |
| --- | --- | --- | --- | --- |
| Cth | 11 | 15.28 ± 9.09 | 0.6876 | 0.4060 |
| shTECRL | 17 | 16.63 ± 8.95 |  |  |

**Table 4.** 1 μM ISO induced faster beating rate of TECRL knockdown hiPSC-CMs versus baseline.

|  | n | bpm (mean±SD) | rate (Hz) | *p* | *t* |
| --- | --- | --- | --- | --- | --- |
| Cth | 11 | 43 ± 10.50 | 0.72 | 0.0044 | 3.143 |
| shTECRL | 17 | 56 ± 8.49 | 0.93 |  |  |

**Table 5.** Baseline of contraction of TECRL overexpression hiPSC-CMs versus baseline.

|  | n | bpm (mean±SD) | rate (Hz) | *p* | *t* |
| --- | --- | --- | --- | --- | --- |
| Cth | 11 | 33.2 ± 9.35 | 0.55 | 0.1861 | 1.365 |
| OeTECRL | 14 | 46.4 ± 28.10 | 0.77 |  |  |

**Table 6.**
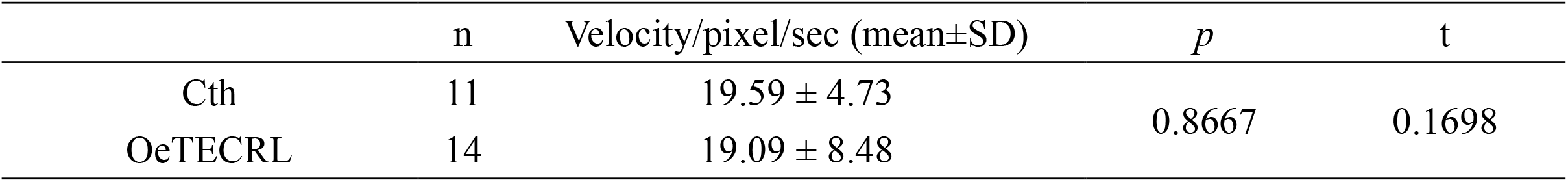
Velocity/pixel/sec of TECRL overexpression hiPSC-CMs versus baseline.

|  | n | Velocity/pixel/sec (mean±SD) | *p* | *t* |
| --- | --- | --- | --- | --- |
| Cth | 11 | 19.59 ± 4.73 | 0.8667 | 0.1698 |
| OeTECRL | 14 | 19.09 ± 8.48 |  |  |

**Table 7.** 1 μM ISO induced faster beating rate of TECRL overexpression hiPSC-CMs versus baseline.

| | n | bpm (mean±SD) | rate (Hz) | *p* | t |
| --- | --- | --- | --- | --- | --- |
| Cth | 11 | 42.0 ± 10.00 | 0.7 | 0.0027 | 3.337 |
| OeTECRL | 14 | 88.5 ± 16.97 | 1.475 | | |

**Fig 6.**
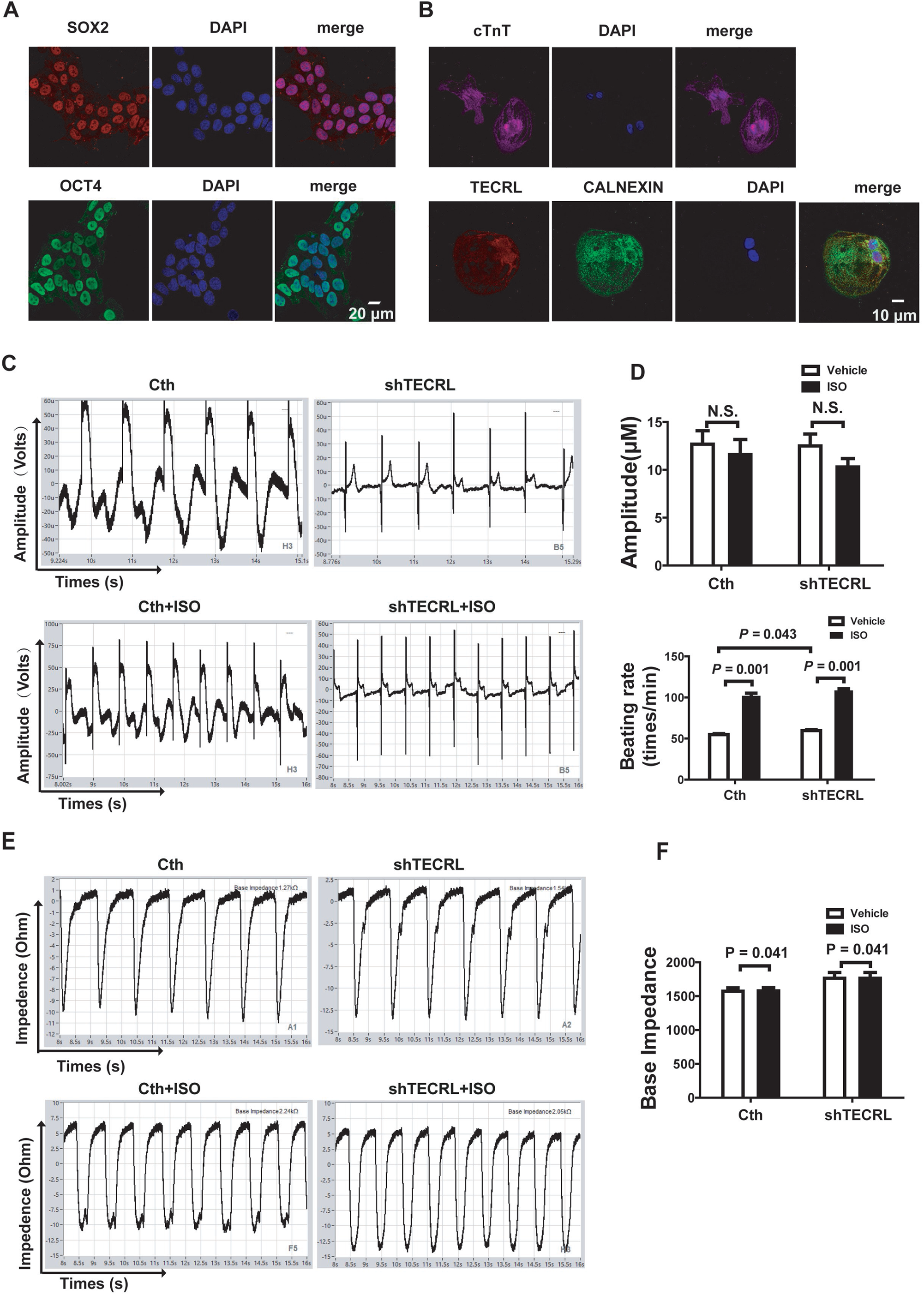
HiPSC-CMs characteristics and the ISO effects on hiPSC-CMs. A. Representative immunofluorescence images of SOX2 and OCT4 in the iPSC (n=3). B. Representative immunofluorescence images of cTnT, TECRL, CALNEXIN in the hiPSC-CMs (n=6). C-D. Representative extracellular field potential(EFP)traces and quantification of the beat rate, and amplitude with Nanion software analysis (n=8). E-F. Representative impedance (IMP) and quantification of base impedance in hiPSC-CMs (n=8). Values are means ± SE. *P* < 0.05 was considered significant.

### 2.6 TECRL deletions disturbed Ca^2+^ regulatory proteins

Strikingly, TECRL knockdown significantly activated P38 phosphorylation in hiPSC-CMs (Fig. 7A), which is consistent with *in vivo* results (Fig. 5B, 5D). As impaired Tecrl signaling increased cell apoptosis (Fig. 2), we found that BAX was also increased, while BCL-2 decreased in hiPSC-CMs (Fig. 7A). *In vitro*, calcium regulating signaling such as CASQ2 was also decreased, while NCX increased in hiPSC-CMs after TECRL knockdown (Fig. 7B). The expression of p-CaMKII was also activated following the TECRL knockdown (Fig. 7C), whereas it was suppressed after TECRL overexpression in the hiPSC-CMs (Fig. 7D).

**Fig 7.**
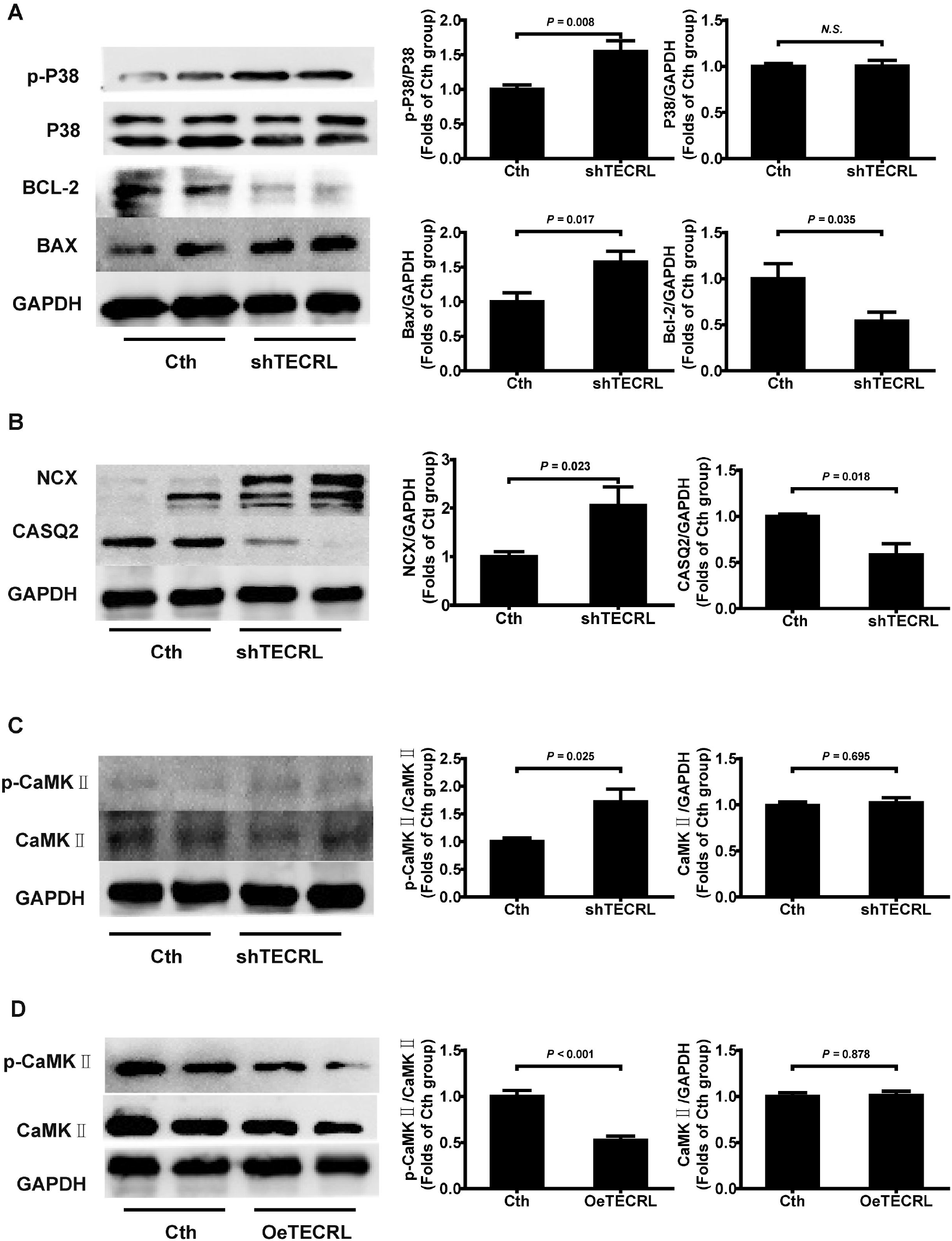
TECRL deletion disturbs calcium regulatory proteins. A. Representative of western blotting images and quantification of p-P38, P38, BAX, and BCL-2 expression in hiPSC-CMs following TECRL knockdown (n=6). B. Representative of western blotting images and quantification of the expression NCX and CASQ2 in the hiPSC-CMs following TECRL knockdown (n=6). C-D. Representative of western blotting images and quantification of p-CaMKII and CaMKII expression in hiPSC-CMs following TECRL knockdown and overexpression (n=6). Values are means ± SE. *P* < 0.05 was considered significant.

### 2.7 CaMKII inhibitions reverses calcium transient abnormalities caused by TECRL deficiency

CaMKII inhibitor, KN93, can suppress TECRL deficiency induced P38 phosphorylation in H9C2 cells (Fig. 8A), and also suppress the activation of CaMKII induced by TECRL deficiency (Fig. 8B). We also found that KN93 could relieve TECRL deficiency induced calcium signaling (NCX, and CASQ2) abnormality (Fig. 8C). Interestingly, TECRL overexpression can also partially reverse the relative expression of p-P38/P38, p-CaMKII / CaMKII, NCX, and CASQ2 in H9C2 cells (Fig. 8D). Furthermore, we found that CaMK II inhibitor, KN93 can partially inhibit cardiomyocytes beating rate (Fig. 9A-B), and increase impedance (Fig. 9C-D), while cardiomyocytes beating amplitude still kept stable (Fig. 9A-B). In a word, KN93 has a crucial role in regulation calcium homeostasis in TECRL deficiency cardiomyocytes.

**Fig 8.**
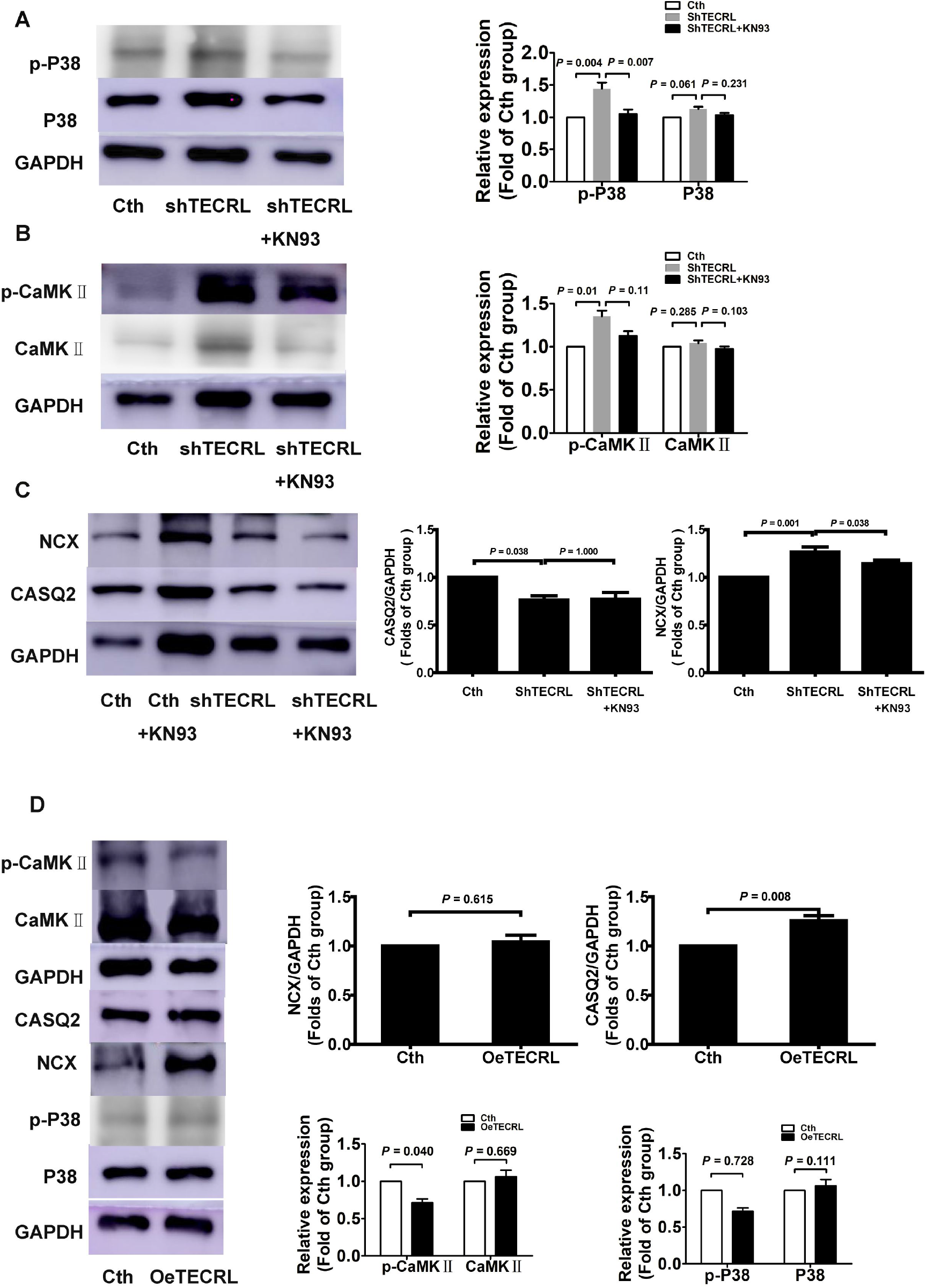
CaMKII inhibitor, KN93 reverses TECRL deletion induced calcium signaling disturbance. A-C. Representative of western blotting images and quantification of p-P38 and P38 (A), p-CaMKII and CaMKII (B), NCX and CASQ2 (C) expression in H9C2 cells upon KN93 pretreatment (n=6). D. Representative of western blotting images and quantification of p-P38, P38, p-CaMKII, CaMKII, NCX, and CASQ2 expression in H9C2 cells following TECRL overexpression (n=6). Values are means ± SE. *P* < 0.05 was considered significant.

**Fig 9.**
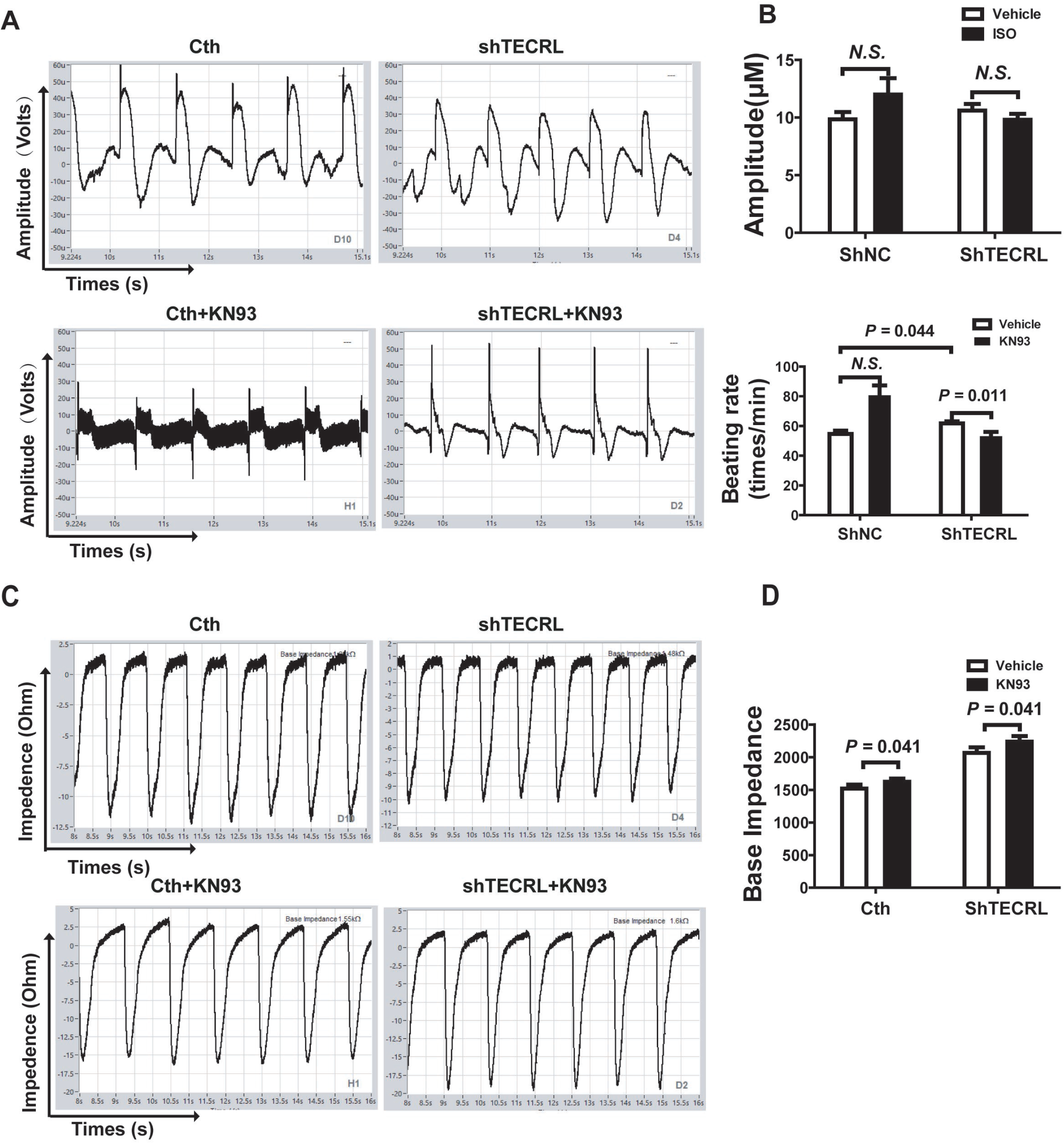
KN93, CaMKII inhibitor effects on hiPSC-CMs function. A-B. Representative images of EFP traces and quantification of beat rate, and amplitude with Nanion software analysis (n=8). C-D. Representative images of IMP and quantification of base impedance in hiPSC-CMs (n=8). Values are means ± SE. *P* < 0.05 was considered significant.

## 3. Discussion and conclusions

In this study, we performed electrophysiological, histological, cellular and molecular characterization of the Tecel KO mice. First, we were successful in generating a model of Tecrl KO mice. There are much more multiple premature ventricular beats and ventricular tachycardia after epinephrine plus caffeine injection in the Tecrl KO mice, indicating that the Tecrl KO mice are sensitized to epi/caffeine induced CPVT phenotypes. Secondly, Tecrl KO mice exhibited much more collagen deposition and disturbed cardiac mitochondria function. Mechanistically, Tecrl deficiency induces overestimation of CaMKII pathways, alters P38MAPK signaling, and decreases the protein stability of RyR2 (one of the crucial calcium regulating). Furthermore, KN93 can partially relieve TECRL deficiency induced cardiomyocytes calcium handling dysfunction. Thus, the Tecrl mouse is a typical model for developing specific therapeutic targets for CPVT and provides a useful tool to shed light on the complex pathogenesis of recessive CPVT.

In last two decades, some CASQ2, RyR2 gene mutations transgenic mice models have been generated. In 2007, Song’s group generated two mutants CASQ2 knock-in mice models (D307H missense mutation CASQ2^(307/307)^ or a CASQ2-null mutation CASQ^(DeltaE9/DeltaE9)^), and both two models developed bidirectional-polymorphic VTs on sympathetic activation (Song et al, 2007). In 2005, a conditional knock-in mouse model carrier of the RyR2^R4496C^ mutation was developed, and this model was aimed to demonstrate that RyR2 mutations can reproduce the arrhythmias observed in CPVT patients for the first time (Song et al, 2007). In 2009, increased Ca^2+^ sensitivity of RyR2 mutation mice (RyR2^R4496C^) underlied CPVT molecular mechanism (Fernandez-Velasco et al, 2009). RyR2^R2474S^ and RyR2^N2386I^ induced CPVT model displayed a mild ER stress response in pancreatic β cells, associated with chronic ER calcium depletion due to RyR2 channel leakage (Santulli et al, 2015). Liu’s group also generated knock-in mice that bear the RyR2^A165D^ mutation (Xiong et al, 2018). All of these indicating that RyR2 and CASQ2 transgenic mice could develop bidirectional and polymorphic VT, which represents extraordinary similarity with the clinical manifestations of CPVT patients. Howerer, there are still no clues to what it would be in a Tecrl mouse model.

It is well known that early clinical features of arrhythmia are mainly mild palpitations, chest tightness, dizziness, and occasionally colic, which can easily be ignored (Stern et al, 2003). Even if the early electrogram shows normal, there is a risk of sudden death in the later stage. The first-line therapy to prevent adrenergically mediated arrhythmias in CPVT is β-blockers (De Ferrari et al, 2015). However, follow-up data in different populations showed recurrence of arrhythmic events in about 25% of patients (Yan et al, 2013), indicating the incomplete protection of this treatment. Devalla’s group first made the connection between TECRL mutation and CPVT (Devalla et al, 2016), and our previous work also confirmed that TECRL mutation (c. 587 C>T and c.918+3T > G) may contribute to CPVT (Xie et al, 2019). In 2020, Webster’s group conducted an international multi-center research and summarized the life-threatening arrhythmias with autosomal recessive TECRL variants (Webster et al, 2020). All of above demonstrating that TECRL was a novel CPVT disease-causing gene. Herein, we developed a mouse Tecrl KO model via CRISPR/Cas9 technique and utilized a combination of electrophysiology approaches and histological analyses to characterize the Tecrl mice model. Upon epinephrine plus caffeine stimulation, multiple premature ventricular beats and VT were observed in the Tecrl KO mice (both at the 4^th^ and 8^th^ weeks) (Fig. 3, and Fig.s2). One of the obvious characteristics of CPVT is calcium handling dysfunction (Bezzerides et al, 2019). We also find that the calcium transient increase remarkedly slowed, accompanied by a decrease of calcium transient amplitude and time to peak (Fig. 4C-D). Thus, all of these electrophysiological tests prompted that Tecrl KO mouse model can faithfully recapitulate CPVT phenotype. Our Tecrl mouse model provides a useful tool to explore early treatment of CPVT and time window for treatment.

By isolating cardiomyothes from the Tecrl KO mice, we found that mitochondria morphology showed some disturbances, such as irregular arrangement, swelling, and vacuolated and disrupted cristae (Fig. 4A). These results are consistent with mitochondrial alterations described in mouse pancreatic islet from RyR2 knockout mice (Santulli et al, 2015). Priori’s group observed that approximately half of cardiomyocytes from the Het mice (RyR2^R4496C^) hearts present mitochondria dysfunction with crista degeneration and an increase in empty cytoplasmic spaces, demonstrating for the first time that RyR2 mutations are associated with mitochondrial damage in the heart (Bongianino et al, 2017). They are also evidenced that RNA-interfered gene therapy reduces mutant RyR2 transcripts and rescues ultrastructural phenotypes, supporting the view of the existence of a causal link between RyR2 dysfunction and mitochondrial abnormalities (Bongianino et al, 2017). Based on the pathological examination, there was obvious fibrosis in the space between myocardial cells (Fig. 1D-E) and much more likelihood of apoptosis (Fig. 2C-D) in the Tecrl KO mice at the age of eight weeks. However, the Tecrl KO mice heart showed normal at the age of 4/5 weeks (Fig. 1-2), indicating that long term Tecrl deficiency induces cardiac dysfunction. It is unclear whether the heart has similar structural changes in patients with CPVT. These findings give another glimpse of calcium regulation affecting mitochondrial function, yet the related mechanisms need to be further explored.

Ca^2+^/calmodulindependent protein kinase II (CaMKII), has been intensively studied for its deleterious roles in experimental heart disease models (Dobrev & Wehrens, 2014; Houser, 2014). Pu’s group recently reported that administration of AAV9-GFP-AIP (CaMKII activates inhibitor) to neonatal mice with a known CPVT mutation (RyR2^R176Q/+^) effectively suppressed ventricular arrhythmias induced by either β-adrenergic stimulation or programmed ventricular pacing, without significant proarrhythmic effect (Bezzerides et al, 2019). Similarly, in our Tecrl KO mouse model, we also found that CaMKII signaling was excessive activation, while the CaMKII signaling was partially abated after TECRL overexpression *in vitro* hiPSC-CMs (Fig. 7). This indicates that TECRL deficency plays crucial roles in regulating CaMKII signaling, leading to CPVT. The SR Ca^2+^ leak not only depends on the properties of diastolic RyR2-closure, but also on SR Ca^2+^ load (Eiringhaus et al, 2019). We find that CaMKII inhibitor, KN93 can suppress TECRL deficiency induced P38 phosphorylation, relieve calcium signaling (NCX, and CASQ2) abnormality (Fig. 8), and inhibit cardiomyocytes beating rate (Fig. 9A-B). Collectively, these findings are consistent with an overlapping LQTS/CPVT phenotype in Tecrl KO mice as well as a disturbed calcium handling and increased propensity for DADs during catecholaminergic stimulation (Devalla et al, 2016).

HiPSC offers opportunities to model cardiac diseases *in vitro* and is especially valuable when myocardial tissue of the patient is not accessible to determine the consequences of a particular mutation. Using hiPSC-CMs, we demonstrated that the cells recapitulate salient features of the disease phenotype based on Matlab analysis and Nanion analysis (Fig. 6, Fig. 9, and Tables 2-7). It was reported that CaMKII activates p38MAPK, which, in turn, activates mitochondrial death pathway by producing an unbalance between the expression of pro- and anti-apoptotic proteins (Bax, Bcl-2)(Baines & Molkentin, 2005; Nguyen et al, 2004). Altered calcium homeostasis is a hallmark of CPVT phenotype, and mutations in RyR2 or CASQ2 have been identified to cause this disease. Devalla’s group reported that protein levels of RyR2 and CASQ2 were significantly downregulated in TECRL Hom-hiPSC-CMs (generated and induced from the CPVT patient)(Baines & Molkentin, 2005; Nguyen et al, 2004). Interestingly, our data showed that p38MAPK signaling were partially increased accompanied with a series of calcium regulating signaling both *in vitro* and *in vivo* (Fig. 5 and Fig. 7). In our work, clinical blood samples from the CPVT families (Xie et al, 2019) were reprogrammed into pluripotent stem cells, which were then divided into cardiomyocytes (results not shown).

In line with our previously CPVT patients clinical information(Xie et al, 2019), cellular functional data from hiPSC-CMs and Tecrl KO mice point toward overlapping features of LQTS and CPVT with triggered activity-mediated arrhythmogenesis. Using Tecrl KO mice, hiPSC-CMs, and H9C2 cell models, we reveal that much more multiple premature ventricular beats and ventricular tachycardia were observed in TECRL deficiency cardiomyocytes upon epinephrine plus caffeine stimulation. Instability of calcium was due to the overactivity of CaMKII pathways, P38MAPK signaling, and the decrease of RyR2 protein stability (one of the crucial calcium signaling). KN93 has a crucial role in regulation calcium homeostasis in TECRL deficiency cardiomyocytes. Our work provides a novel CPVT murine model and provides support for targeting TECRL in treating CPVT. Further studies are needed to elucidate the exact mechanisms underlying the electrical phenotype, to assess the prevalence of TECRL mutations in CPVT, and to reveal the cause of unexplained cardiac arrest.

### Short coming

From the CPVT family clinical blood samples(Xie et al, 2019), we found that the *TECRL* protein level of the patient was much lower than his parents’ (Fig. s1 C), indicating that a Tecrl KO mouse model can mimic clinical phenotypes of CPVT disease. However, we still need to construct the point mutation mouse and further test them for arrhythmias.

## 4. Materials and methods

### 4.1 Animal experimentation

WT and Tecrl KO mice at the age between 4 and 8 weeks old were used for this study. They were raised under controlled conditions (12 h dark-light cycles, 22±2 °C temperature, and 45-55% relative humidity) with a maximum housing capacity of five mice per cage, where unrestricted access to diet and water was provided. All compounds, such as epinephrine, ISO, caffeine, were dissolved in DI water. All animal studies were subject to the approval of the Ethics Committee of Experimental Research of Shanghai Children’s Hospital, Shanghai Jiaotong University.

### 4.2 CRISPR/Cas9-mediated Tecrl KO mice

Tecrl KO mice were generated by Biocytogen (Beijing, China) with the EGE system based CRISPR/Cas9 technology. Designed by the CRISPR design tool (http://www.sanger.ac.uk/htgt/wge/), two single-guided RNAs (sgRNAs) targeted upstream of exon 3 and downstream of exon 4 of Tecrl, respectively.

### 4.3 Morphological and histological analyses

Heart tissues were excised, fixed in 10% formalin, and embedded in paraffin. Heart sections (4 μm) were stained with Masson and hematoxylin and eosin (HE) (Sigma, Germany) staining, according to the manufacturer’s instructions. Pathological changes were observed under an optical microscope (Olympus, Japan).

### 4.4 Immunofluorescent and TUNEL staining

The apoptosis in heart tissue was measured through a TUNEL assay according to the manual. In brief, cardiac tissues embedded with optimal cutting temperature compound (OCT) were cryosectioned at a thickness of 4-7 μm. The slices were then balanced at room temperature for 30 min and incubated with the TUNEL reaction mixture in the dark for 60 min. Ten fields were randomly selected for each section, and TUNEL-positive cells were counted and averaged. All histological examinations were performed in a blinded fashion.

### 4.5 Surface electrocardiogram

Animals were anesthetized through intraperitoneal injection of 100 mg/kg pentobarbital and placed on a heating pad. Respiratory rate and loss of toe-press reflex were used to monitor the level of anesthesia(Hwang et al, 2014; Watanabe et al, 2009). Electrocardiogram (ECG) was performed on mice at the 4th and 8th weeks with a non-invasive small animal electrocardiogram (INDUS Technology, Inc, CA, USA). Baseline ECG was recorded for 5 min, followed by an additional 15 min after intraperitoneal administration of epinephrine (2 mg/kg) plus caffeine (120 mg/kg). Quantification of arrhythmic episodes was performed offline by 2 blinded investigators.

### 4.6 Transmission electron microscopy

Transmission electron microscopy (TEM, Thermo Fisher, MA, USA) for morphological analysis was performed at Shanghai Institute Precision Medicine, Ninth People’s Hospital, Shanghai Jiaotong University School of Medicine. For TEM sample preparation, acute isolated cardiomyocytes were first fixed with ice-cold 2.5% glutaraldehyde at 4°C overnight. Ultrathin sections were stained with uranyl acetate and lead citrate. The resulting sample with 90–100 nm thickness was mounted on a 200-mesh copper grid and imprinted using an FEI Titan Krios G3 Spirit transmission electron microscope (Thermo Fisher, MA, USA). In each sample, 6-8 visual fields were randomly selected, and then the number of mitochondria was counted through image J software. All analyses were performed blind to the observer.

### 4.7 Isolation and culture of adult cardiomyocytes

Adult mice cardiomyocytes were isolated from the WT and Tecrl KO mice following the published protocol(Judd et al, 2016; Liu et al, 2019). Briefly, a mouse was injected with 150 or 300 units of heparin for 20 min. The mouse was anesthetized by intraperitoneal injection of 100 mg/kg pentobarbital. The heart with lungs attached was quickly removed and put into the ice-cold buffer. Removed the lung, uncovered the ascending aorta, cannulated via the aorta, tied the aorta and mounted on a modified Langendorff perfusion system. Subsequently, the heart was perfused with 37 °C oxygenated Krebs-Henseleit buffer solution minus calcium, and then perfused with enzyme solution, which contains 1.17 mg/ml Collagenase Type II (Thermo Fisher, MA, USA) and 1.67 mg/ml Collagenase Type IV (Sigma, Germany) for 30 min. Afterward, the heart was cut into small pieces for further digestion under gentle agitation in the enzyme solution. Rod-shaped adult cardiomyocytes were collected by centrifugation and followed by the gradual addition of CaCl_2_. Freshly isolated cardiomyocytes were plated on 25-mm cover-slips pre-coated with 40 μg/ml laminin for 1 h (Thermo Fisher, MA, USA). All cardiomyocytes were cultured in M199 medium (Sigma, Germany) supplemented with 10% fetal bovine serum (Gibco, USA) at 37 °C and 5% CO_2_ in a humidified incubator. Only excitable, rod-shaped, and quiescent cells were used for further studies.

### 4.8 Ca^2+^ transients and cell shortening in canine cardiomyocytes

Cell shortening and intracellular calcium concentrations were measured as described previously(Oda et al, 2005). In brief, cardiomyocytes were incubated with 5 μmol/L Fura-2 AM, in DMEM culture medium (Gibco, USA) containing 10% fetal bovine serum (Gibco, USA) for 30 min, then washed twice to remove excessive unbound Fura-2 AM. Cells were stimulated by a field electric stimulator (IonOptix, Milton, Massachusetts, USA) at 1 Hz. A dual excitation spectrofluorometer was used to record fluorescence emissions (at 505 nm) under the exciting wavelength of 340 or 380 nm. Intracellular calcium was monitored as the ratio of the fluorescence intensity of the cell at 340 excitation and that at 380 nm.

### 4.9 Human induced pluripotent stem cell-based models and virus infection

A healthy volunteer was consented to participate in this study, following the protocol approved by the Shanghai Children’s Hospital Institutional Review Board. Peripheral blood mononuclear cells were reprogrammed to pluripotency using the CytoTune Sendai reprogramming kit (Thermo Fisher, MA, USA). These colonies were then stained for the pluripotency markers SOX2, OCT4. Human induced pluripotent stem cells (hiPSCs) were differentiated to human induced pluripotent stem cell-derived cardiomyocytes (hiPSC-CMs) according to published protocol(Bezzerides et al, 2019). All of the hiPSCs were maintained in nutristem medium (Biological Industries, Israel) and passaged in accutase enzyme cell detachment medium (Gibco, USA) every three to five days. Culture dishes were pre-coated with Matrigal (BD, USA), diluted at 1:400. After at least 20 to 30 passages, hiPSCs were seeded to differentiation to hiPSC-CMs based on the timeline. On day 1-2, Chir (4-6 μM) was added to aliquot of media 1. Day 3-4, IWR (5 μM) was added to aliquot of media 1. Day 5-6, the media in wells was replaced with media 1 with no chemicals added. Day 7-10, the media in wells was replaced with media 3. Day 11-13, the media in wells was replaced with media 2. During day 10 through day 15, contraction should be visible under the microscope, indicating the differentiation is done successfully. Between day 14 and day 15, the cells need to be placed onto a fresh plate to remove fibroblast and other cell types that were generated along with cardiomyocytes differentiation.

A DNA fragment containing full-length Tecrl cDNA was obtained through PCR amplification. The enzyme restriction sites and the full length of TECRL were inserted into PGMLV-CMV-MCS-3×Flag-PGK-Puro plasmid to construct the recombinant vector. The recombinant plasmid and lenti-virus-packaged helper plasmid were transferred to 293T cells to collect lenti-viruses.

### 4.10 Cardiomyocytes contractility assay

An Olympus IX83 inverted microscope (Olympus, Japan) was used to record five to fifteen-seconds of video imaging on the clusters of 2 to 4 hiPSC-CMs maintained at 37 °C and 5% CO_2_. Contraction of cardiomyocytes was acquired with differential interference contrast optics (100X, 60 frames/s) at specific time points. We conducted cell pacing at 1 Hz while spontaneously beating. To quantify velocity of contraction, we used velocity/pixel/sec with Matlab software (Mathworks, Natick, MA) to measure contractility of cardiomyocytes. According to the peak differences between frames in each absolute pixel, the onset of contraction of each video was identified. Velocity/pixel/sec was acquired *via* diving displacement frame by frame, then converting vector displacements into velocities(Li et al, 2014).

### 4.11 CardioExcyte 96 recordings

The CardioExcyte 96 (Nanion Technologies GmbH, Germany) was used in impedance and extracellular field potential (EFP) mode. The hiPSC-CMs were seeded on the sensor plate that were mounted on the CardioExcyte 96 with an incubation chamber (temperature 37 °C, 5% CO_2_). The stimulation was 1 Hz. Amplitude and beat rate were monitored with the CardioExcyteControl software.

### 4.12 Immunofluorescence analysis

HiPSCs and hiPSC-CMs were fixed with 4% paraformaldehyde for 15 min. To eliminate the non-specific binding of antibodies, samples were incubated in tris-buffered saline tween-20 (TBST) containing 3% bovine serum albumin and 0.1% Triton X-100 for 60 min. After blocking, cells were stained with SOX2, OCT4, CALNEXIN, and cTnT antibody (Cell Signaling Technology, USA) at a dilution of 1:100 overnight at 4°C. After rinsing with TBST buffer, the sample was incubated with secondary antibodies conjugated to Alexa Fluor 594 or Alexa Fluor 633 (Invitrogen, USA) at room temperature for 1 h, followed by DAPI staining. Images were acquired with a laser confocal microscope (Zeiss LSM710, Germany).

### 4.13 Translation inhibition

A translation inhibitor cycloheximide (CHX) was used to detect the RyR2 translation. Briefly, the constructed control and shTECRL lentivirus were transiently transfected into H9C2 cells then treated with CHX (500 μM) for 0, 4, 8, and 24 h. Cell proteins were extracted and analyzed by western blotting.

### 4.14 Cell lysate and western blotting

To acquire cell protein for immunoblotting analysis, hiPSC-CMs and H9C2 cells were washed twice with phosphate-buffered saline (PBS) and incubated with cell lysate buffer at 4°C for 5 min while rocking gently. Proteins were extracted from left ventricular myocardial tissue and lysed with cell lysate buffer. The cell lysate buffer contained RIPA, protease inhibitors, and phosphatase inhibitors. Proteins were resolved by SDS-PAGE and transferred onto polyvinylidene fluoride membrane (Millipore, Bedford, MA, USA). The membranes were incubated with a primary antibody against TECRL (Invitrogen, USA), p-P38, P38, Bax, Bcl-2, CASQ2, Ncx, Ryr2, p-CaMKII and CaMKII (Cell Signaling Technology, USA) at 1:1000 dilution with the blocking buffer. Membranes were incubated with a secondary antibody conjugated horseradish peroxidase (HRP) (Cell Signaling Technology, USA) in the blocking buffer at 1: 2000. After washing, membrane was visualized by electro-chemiluminescent (ECL) analysis. Densities of immunoblot bands were analyzed using a scanning densitometer (model GS-800, Bio-Rad Laboratories, Hercules, CA, USA) coupled with Bio-Rad personal computer analysis software.

### 4.15 Statistical analysis

Differential expression analysis for any two groups was performed using the DESeq2 R package (1.26.0). A *P* value < 0.05 and fold Change ≥ 2 was set as the threshold for significantly differential expression. Results are expressed as mean ± SEM. Statistical analysis was performed using SPSS software, version 21.0 (SPSS, Inc., USA). Comparisons among groups were performed by one-way ANOVA. Paired data were evaluated by two-tailed Student’s t-test. Statistical significance was considered when *P* < 0.05.

## Acknowledgments

This work was supported by National Natural Science Foundation of China (NSFC) (81900437, 82170518), the Shanghai Jiaotong University Medical Technology Crossing Project (YG2021ZD26), the Shanghai Science and Technology Committee (19411963600), and Shanghai Children’s Hospital (2019YQ006). No benefit in any form has been or will be received from a commercial organization directly or indirectly.

## The paper explained

### Problem

Mutations in the genes for RyR2 and CASQ2, two mainly subtypes of CPVT, have been identified. However, the TECRL gene mutation was rarely reported, the TECRL protein functions in CPVT or other cardiovascular disease are still not well understood.

### Results

Tecrl KO mice showed much more multiple premature ventricular beats and ventricular tachycardia, after epinephrine plus caffeine injection, which can mimic CPVT phenotypes.

Intracellular calcium amplitude was reduced, while time to baseline of 50 was increased in acute isolated cardiomyocytes upon 1-Hz electrical stimulation, accompanied with a decrease of calcium regulating signaling, such as CASQ2 or RyR2.

Overexpression of TECRL and KN93 can partially reverse cardiomyocytes calcium dysfunction, regulation of the calcium homeostasis and this is p-CaMKII/CaMKII dependent.

### Impact

A new CPVT model was constructed, which can simulate CPVT clinical phenotypes, such as pleomorphic ventricular velocity. The Tecrl mouse is a typical model for developing specific therapeutic targets for CPVT and provides a useful tool to shed light on the complex pathogenesis of recessive CPVT.

## Author contributions

Cuilan Hou, Lijian Xie, and Tingting Xiao designed and operated the project. Xunwei Jiang, Yongwei Zhang, Meng Xu, and Qingzhu Qiu provided the ECG and echocardiography analysis. Cuilan Hou wrote the manuscript with input from Junmin Zheng, Shujia Lin, and Shun Chen. All authors read and approved the final manuscript.

## Supporting information

**Fig s1.**
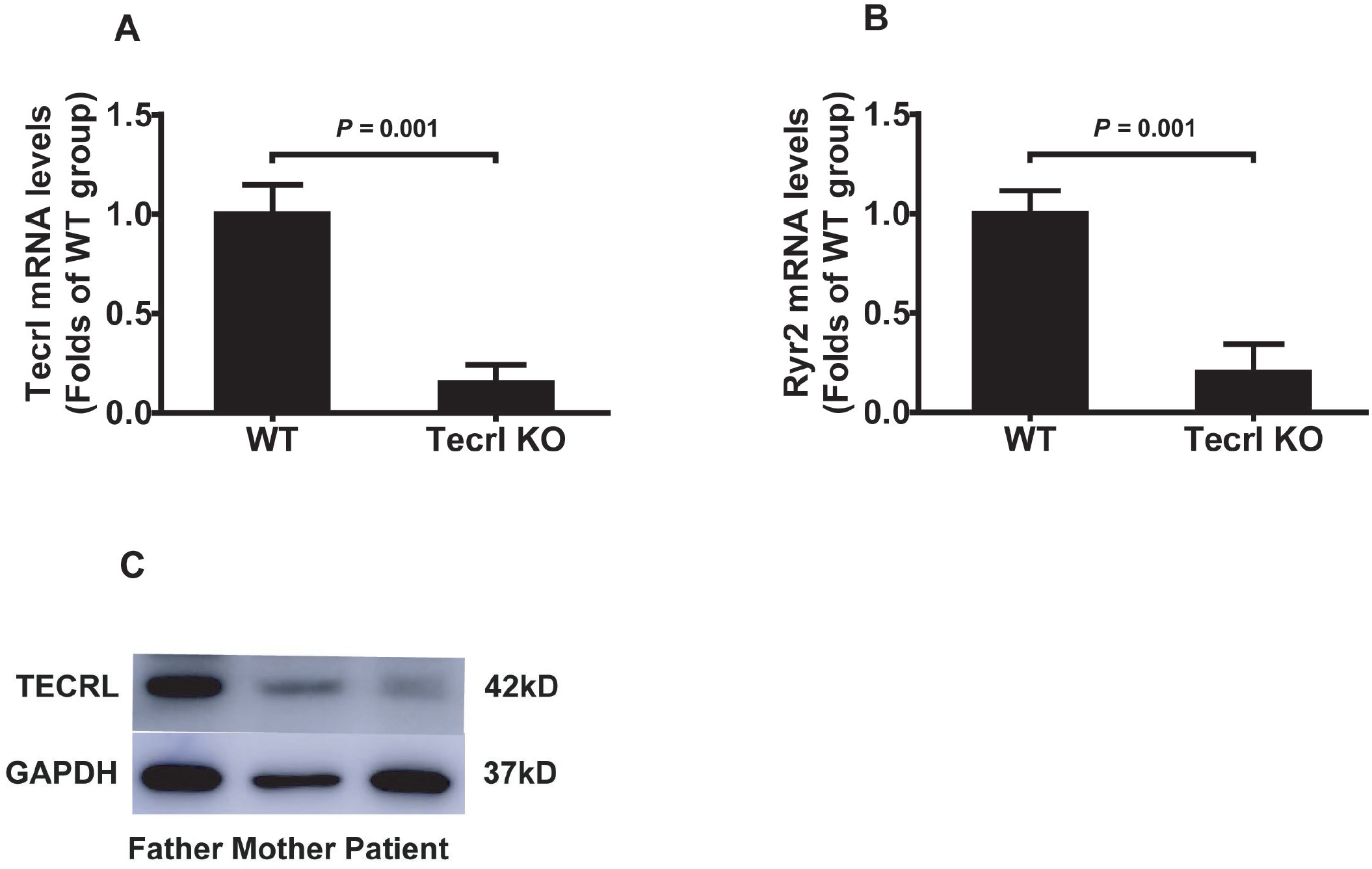
The relative Tecrl expression in mice and CPVT family clinical samples. A-B. Quantification of mRNA levels of Tecrl and Ryr2 in the WT and Tecrl KO mice (n = 6). C. Representative of western blotting images of the patient and his parents (n=1). Values are means ± SE. *P* < 0.05 was considered significant.

**Fig s2.**
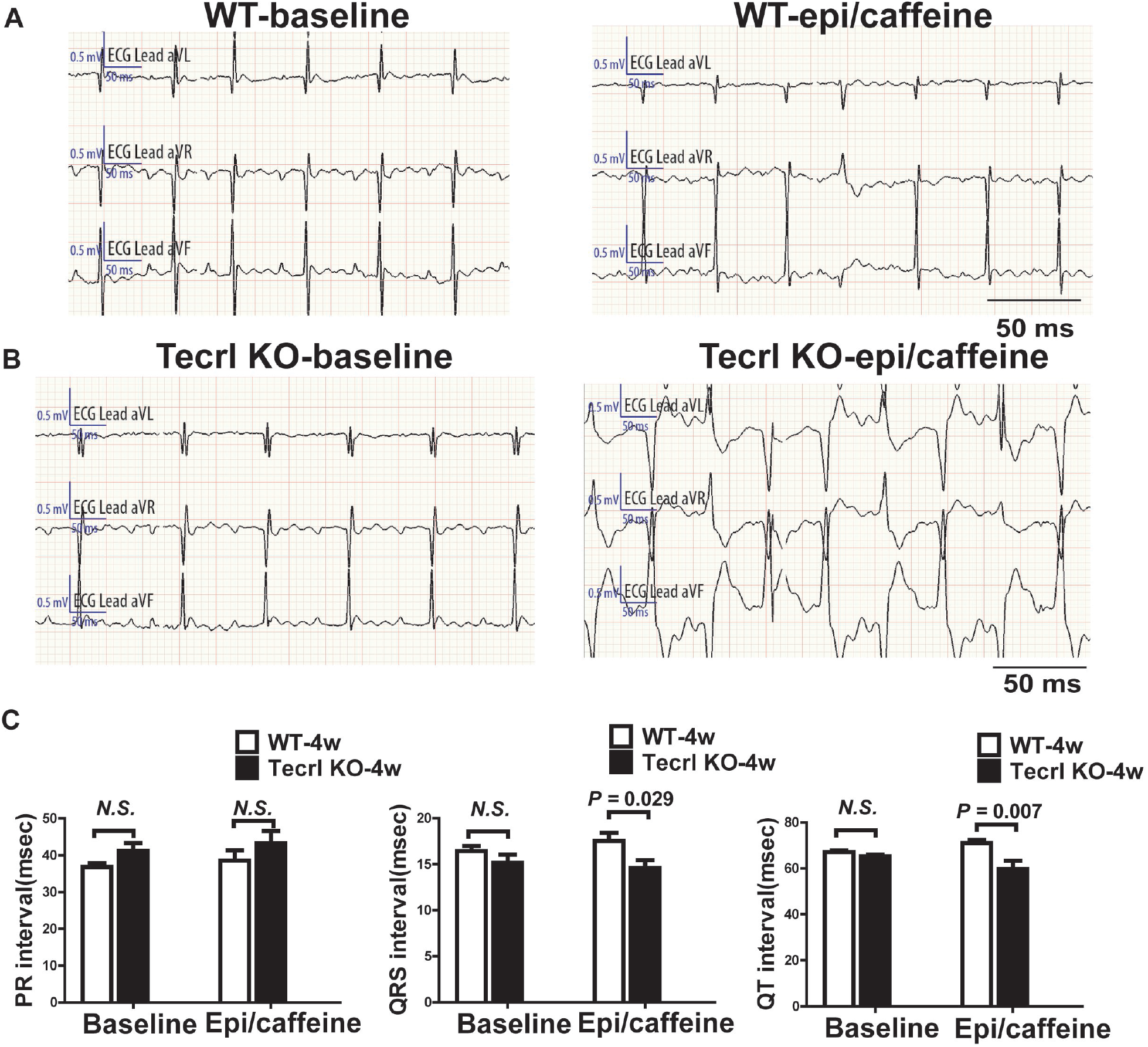
Arrhythmia inductions in the Tecrl KO mice of four weeks. A-B. Representative images of ECG recordings in the WT and Tecrl KO mice before and after epi/caffeine stimulation (n=6). C-D. Quantification of sustained bigeminy, sustained VT, and VT episode in the WT and Tecrl KO mice (n=6). Values are means ± SE. *P* < 0.05 was considered significant.

**Fig s3.**
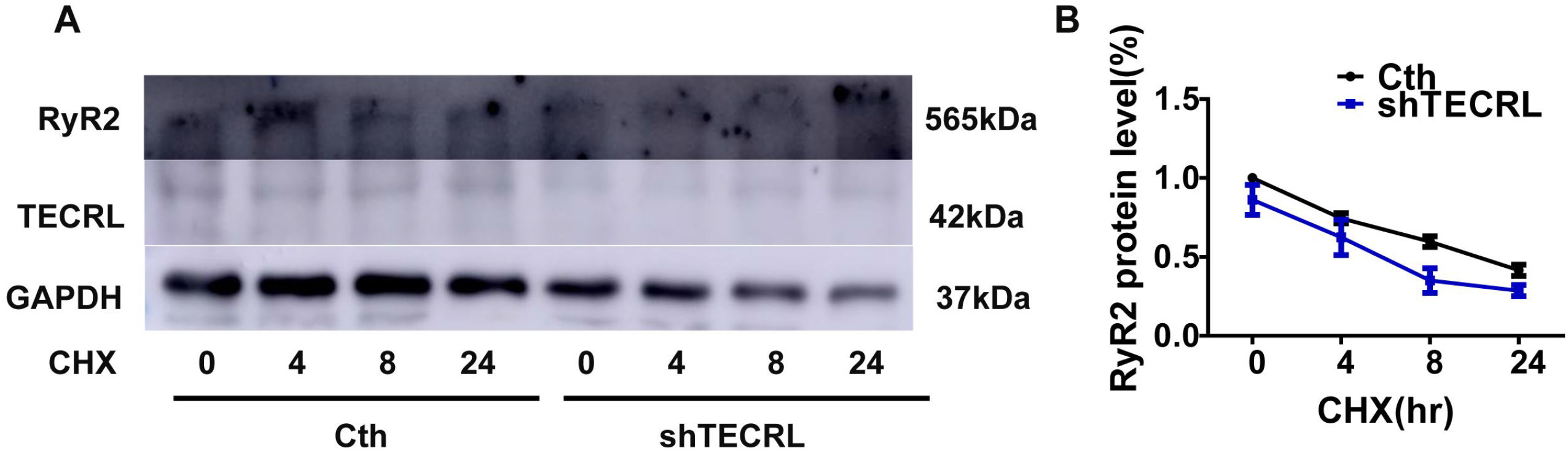
Effects of CHX on RyR2 expression. A. Effects of CHX on RyR2 expression in transfected cells. Immunoblot of RyR2 protein levels between the Cth and shTECRL group after incubation with CHX (500 μM) for 0, 4, 8, and 24 h (n=3). (B) Quantification of relative RyR2 protein levels in transfected cells (n=3). Values are means ± SE. *P* < 0.05 was considered significant.

**Fig s4.**
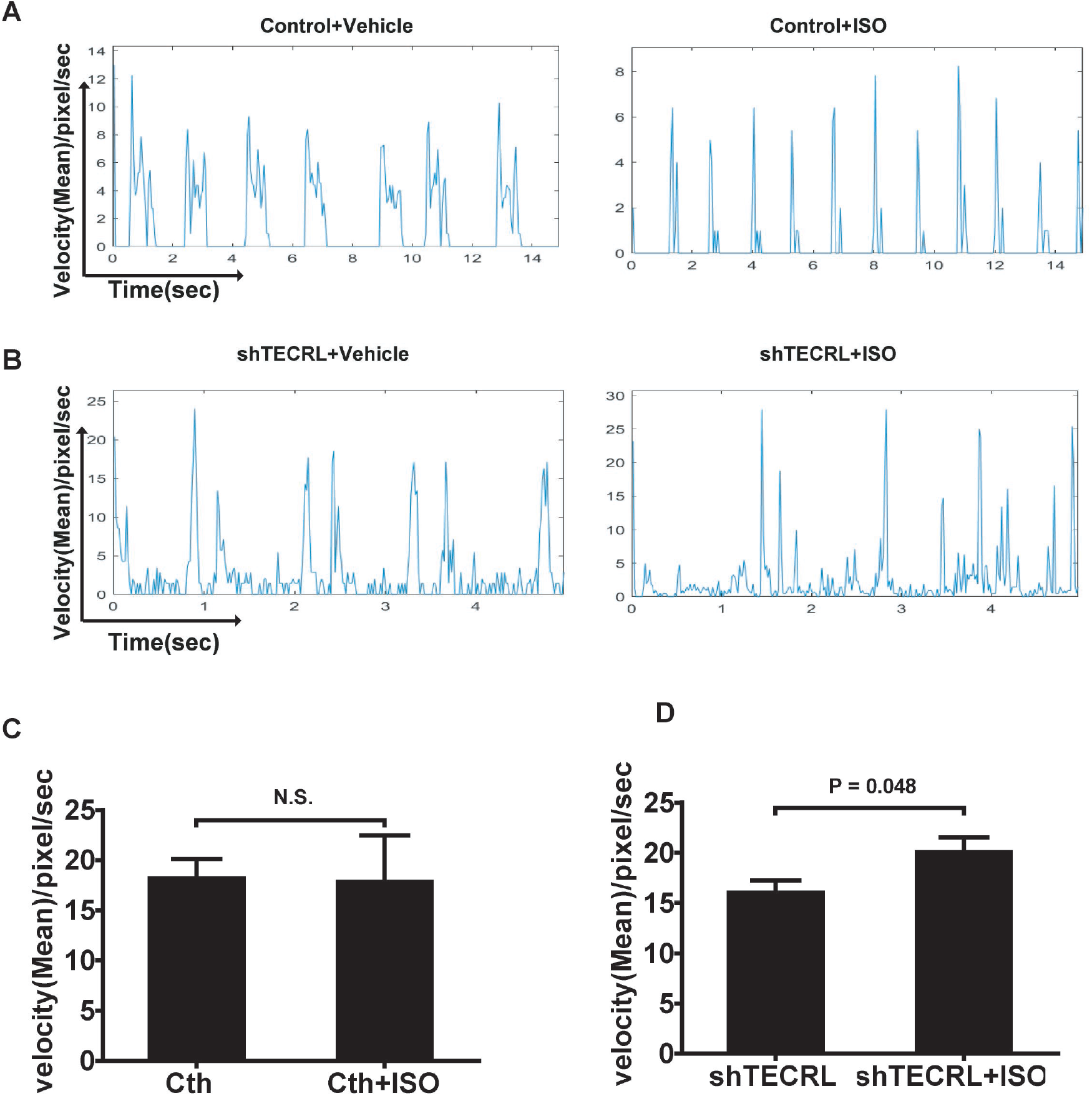
TECRL deficiency effects on cardiomyocytes function. A-D. Representitive absolute amplitude, rhythm and quantification of contraction peak value (velocity (Mean)/pixel/sec) in hiPSC-CMs upon TECRL knockdown and ISO stimulation (n=11). Values are means ± SE. *P* < 0.05 was considered significant.

**Table s1.**
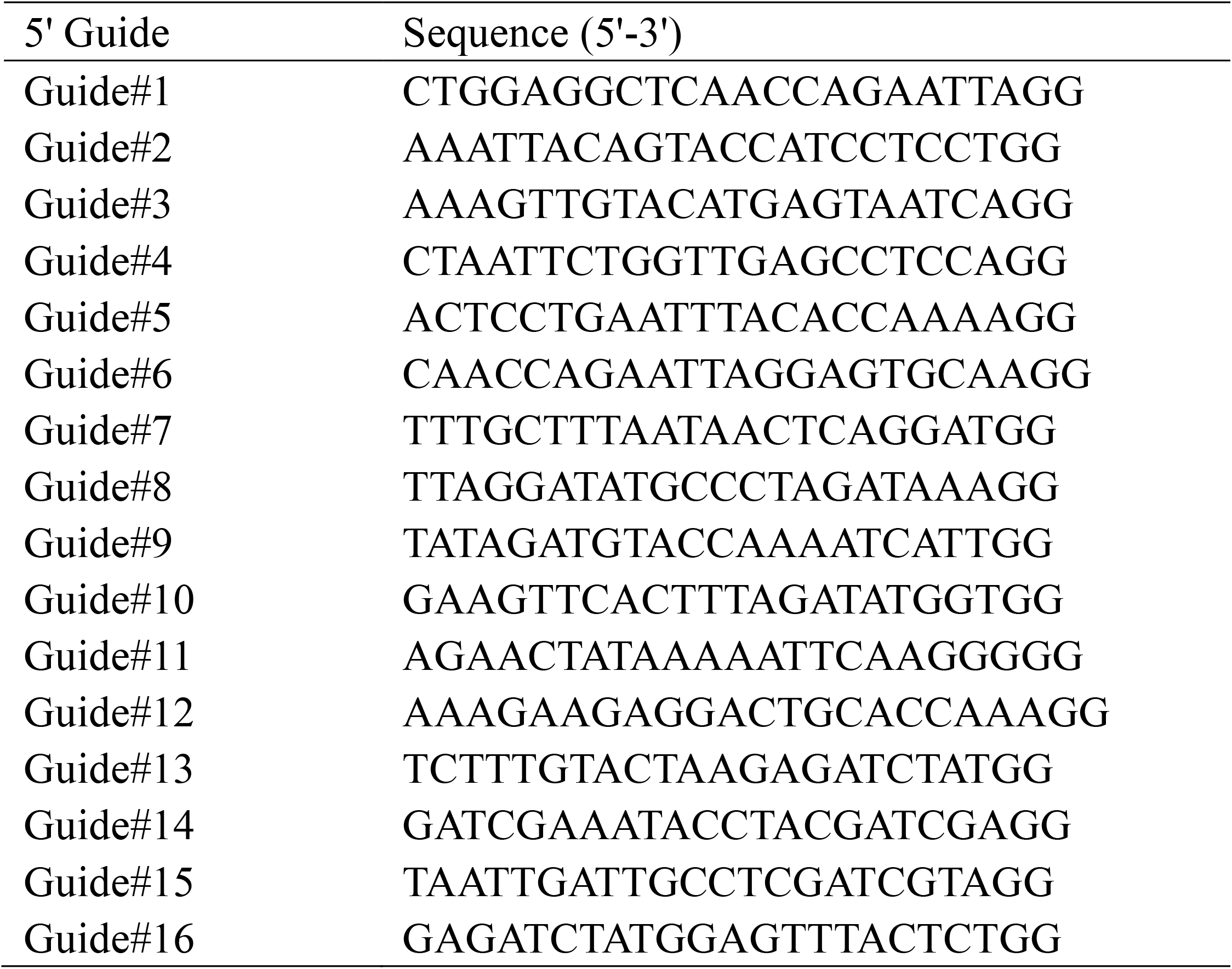
The sequences of sgRNAs.

